# Development and Characterization of the Mode-of-Action of Inhibitory and Agonist Peptides Targeting the Voltage-Gated Sodium Channel SCN1B/β1 Subunit

**DOI:** 10.1101/2023.10.19.562974

**Authors:** Zachary J. Williams, Anita Alvarez-Laviada, Daniel Hoagland, L. Jane Jourdan, Steven Poelzing, Julia Gorelik, Robert G. Gourdie

## Abstract

Treatment of cardiac arrhythmias by targeting ion channels is challenging, with safe and effective therapies remaining an unmet clinical need. Modeling and experimental studies have shown that a voltage-gated sodium channel (VGSC)-rich nanodomain at edge of the gap junction (GJ) called the perinexus could provide new mechanistic insights into normal and abnormal conduction of action potentials in the heart. We have reported that a 19 amino acid SCN1B (β1/β1B) mimetic peptide derived from the immunoglobulin domain of the VGSC subunit called βadp1 acutely disrupts β1-mediated adhesive interactions at cardiac perinexii, prompting arrhythmogenic changes during time courses of up to an hour. In the present study, we sought to gain further insight on βadp1 mode-of-action, as well as identifying new SCN1B (β1/β1B) mimetic peptides, with potential for inhibiting and/or promoting β1-mediated adhesion. This included studies of the effect of βadp1 and related peptides on SCN1B (β1/β1B) Regulated Intramembrane Proteolysis (RIP) - a signaling pathway that has been shown to effect gene transcription, including that of VGSC subunits. Using patch clamp to assay cell-cell contact-associated VGSC activity in cardiomyocytes, and electric cell substrate impedance sensing (ECIS) to assess intercellular adhesion in cells heterologously expressing β1, we find that inhibitory effects of βadp1 can persist for up to 5 hours. However, this acute inhibition is not sustained, with βadp1 effects on β1-mediated adhesion lost after 24 hours. We also determined that a short peptide (LQLEED) near the carboxyl-terminal portion of βadp1 inhibited adhesion in β1-expressing cells in a manner similar to βadp1. Paradoxically, dimeric peptides incorporating a repeat of the LQLEED sequence promoted intercellular adhesion at all time points studied over a 2-day time course. Inhibitory and agonistic peptides were found to effect β1 RIP, with βadp1increasing RIP continuously over 48 hours, whilst dimeric agonists acutely increased RIP at 6 hours post-treatment, but not thereafter. In the presence of DAPT, an inhibitor of RIP, the effects of βadp1 on ECIS-measured intercellular adhesion were lost, suggesting a relationship between RIP and inhibitory effects of the peptide. In sum, we identify novel SCN1B (β1/β1B) mimetic peptides with potential to inhibit and promote intercellular β1-mediated adhesion, possibly including by effects on β1 RIP, suggesting paths to development of anti-arrhythmic drugs targeting the perinexus.

## INTRODUCTION

The United States experiences between 180,000 and 450,000 sudden cardiac deaths annually, with cardiac arrhythmias playing a significant role [1–3]. Many drugs aimed at preventing arrhythmias target ion channels, including the sodium channel Nav1.5, but often have severe side effects [4]. For example, the Cardiac Arrhythmia Suppression Trial (CAST) was successful in reducing arrhythmogenic triggers, but unfortunately the therapies tested were also found to result in an increase in total deaths [5, 6]. This problematic clinical trial record, together with high costs, and the inherent deadly risks associated with targeting heart rhythm disorders, has slowed the development of anti-arrhythmic drugs. While there are new and growing methods of treating arrhythmias, such as catheter ablation [7, 8] neuroscientific therapies [9], or even optogenetic methods [10], there remains considerable academic and clinical interest in new mechanistic targets to address the growing unmet clinical need for safe and effective pharmacologic treatment of disorders of cardiac electrical rhythm [11, 12].

The voltage-gated sodium channel (VGSC) incorporates both alpha(α)-(e.g. Nav1.5) and beta (β)-subunits. VGSC β subunits are encoded by SCN1B-SCN4B (β1-β4) and an alternatively spliced variant of SCN1B (β1B) [13–20]. VGSC β-subunits have been reported to regulate channel excitability by altering channel gating, voltage-dependence of activation and inactivation, and inactivation speed. Additionally, β subunits have assignments in cell adhesion - mediated by an extracellular immunoglobulin (Ig) domain that shares homology with other adhesion molecules [13]. Consistent with adhesive function, SCN1B (β1/β1B) is prominently localized at zones of electro-mechanical contact between cardiomyocytes [21, 22], in particular at the perinexus, an intercalated disc nanodomain adjacent to the gap junction (GJ), co-locating with Nav1.5 [21, 22], other ion channels and scaffolding proteins [23–25].

A growing body of modeling and experimental data indicate that the perinexus has roles in normal and arrhythmogenic conduction of cardiac electrical excitation [21, 26–32]. The VGSC-rich perinexal domain at the GJ edge forms a 20-30 nm wide cleft of extracellular space. Adhesive interactions between Ig domains of SCN1B molecules on cells apposed at the GJ-adjacent cleft have been suggested to contribute to perinexal stability [21]. Two recent studies have provided further insights. First, modeling work by Ivanovic and Kucera showed that the narrow intercellular spacing maintained at the perinexus, together with its adjacency to GJs, profoundly affected extracellular potential dynamics and local patterns of current flow, orchestrating propagation of action potentials via an ephaptic mechanism [26]. Second, Adams et al. demonstrated that the width of the perinexus, and its effect on the ephaptic mechanism, is a key determinant of normal cardiac conduction and not just a phenomenon of pathological states, as had been proposed by some [27]. Consistent with roles in health and disease, increases in cell-cell spacing at the perinexus has been linked to atrial arrhythmia in humans, suggesting that regulation of perinexal width and/or SCN1B (β1/β1B)-mediated adhesion may be targets for treating electrical disturbance to the heart [21, 33]. Further supporting this hypothesis, SCN1B KO mice show widened perinexii. Mutations in SCN1B in humans, including in the Ig domain, have been implicated in several arrhythmia-associated pathologies, including Brugada syndrome and Long QT syndrome [21, 34, 35].

In a previous report, we showed that a SCN1B (β1/β1B) mimetic peptide derived from its immunoglobulin (Ig) domain called βadp1 acutely disrupted β1-mediated function. Over time courses of an hour or less, βadp1 reduced adhesion in cells expressing SCN1B (β1), caused de-adhesion and expansion of perinexii in isolated guinea pig hearts and prompted arrhythmogenic changes in cardiomyocytes and hearts, including loss of sodium channel activity and conduction slowing [21]. In the present study, we investigated short- and long-term effects of βadp1 and related peptides – including novel β1-targeting inhibitory and agonist peptides. We find that the acute effects of β1-targeting peptides, including loss of junctional sodium currents in cardiomyocytes and reductions in adhesion between β1-expressing cells, are maintained for up to 5 hours. It was further determined that over longer periods the response of cells to the peptide altered, with treatment being associated with increased adhesion. Moreover, we show that this increase correlates with up-regulation of Regulated Intramembrane Proteolysis (RIP) of β1. β1 was shown to be modified through RIP almost two decades ago [36]. Recent studies have solidified the importance of β1 RIP by showing that the intracellular domain of β1 translocates to the nucleus and alters transcription of various genes involved in the propagation of electrical activity in the heart, including VGSC complex proteins [37, 38]. The results described herein suggest novel pro-drugs and paths to pharmacologically addressing the VGSC complex, with relevance to potential targeting of the perinexus and its assignments in cardiac conduction.

## RESULTS

### βadp1 Reduces Junctional Sodium Channel Activity, β1 Levels and Adhesion Over an Acute Time-Course

We have previously shown that a 30 minute treatment with βadp1, a 19 amino acid peptide mimetic of the SCN1B/β1/β1B immunoglobulin (Ig) domain, reduces peak sodium channel current (I_Na_) at cell-to-cell junctional contacts between neonatal rat ventricular myocytes (NRVMs) (Fig. 1A) [21]. βadp1 had no effect on sodium channel activity in non-junctional membranes distal from cell-cell contacts, or on whole cell I_Na_, over similar time courses. We used scanning ion conductance microscopy (SICM) –guided smart patch clamp in NRVM cultures to investigate junctional sodium currents over a longer period (Fig. 1A). Similar to what was observed in the earlier study, 50 μM βadp1 reduced junctional I_Na_ significantly after 30 minutes (Fig 1B). However, longer exposure to βadp1 prompted further reductions of junctional I_Na_, reaching levels ∼ 1/3 that of untreated cells after 60 minutes, thereafter leveling off to similar reductions in activity at 90 and 120 minutes (Fig 1B). No change in I_Na_ occurred at junctional contacts in response to scrambled control peptide or in non-junctional membranes in response to βadp1 or control peptide (Fig. 1C). The level of sodium channel activity in non-junctional NRVM membranes was similar to the suppressed levels of I_Na_ measured after 60 minutes or more of exposure to βadp1 in junctional contacts (Fig 1C).

**Figure 1.**
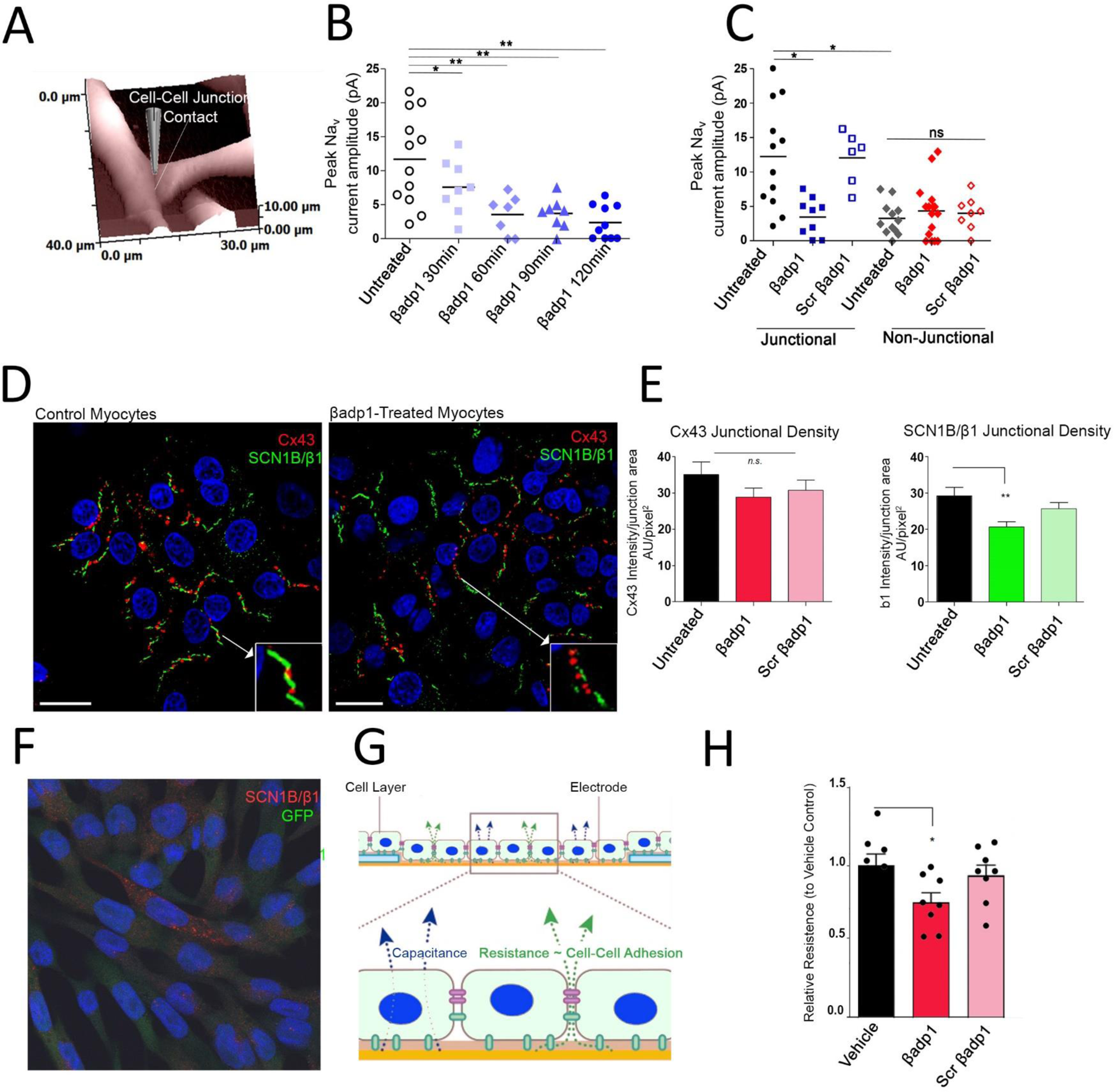
Acute effects of βadp1 on I_Na_, β1 density and intercellular adhesion: **A)** Surface of neonatal rat ventricular myocytes (NRVMs) at a cell-to-cell contact imaged by Scanning Impedance Conductance Microscopy (SICM). A cartoon of a patch pipette is shown near a cell-to-cell contact. **B)** Summary plot of I_Na_ from junctional sites over a 120 minute time course following treatment with 50 μM βadp1 (Untreated Control: n = 12; βadp1 30 minutes: n = 8; βadp1 60 minutes: n = 7; βadp1 90 minutes: n = 8; βadp1 120 minutes: n = 10) **C)** Summary plot of I_Na_ from junctional and non-junctional sites in presence or absence βadp1 or scrambled βadp1 control peptide (50 μM) after 60 minutes (Junctional untreated Control: n = 12; Junctional βadp1: n = 8; Junctional scrambled βadp1: n = 6; Non-Junctional untreated Control: n = 10; Non-Junctional βadp1: n = 12; Non-Junctional scrambled βadp1: n = 6) **D)** Representative confocal images of NRVMs immunolabeled for Cx43 (red) and β1 (green) of control NRVMs or after treatment with βadp1 50 μM for 60 minutes. Insets show magnified views of junctional Cx43 (red) and β1 (green) signals. **E)** Quantification of immunolabeling density of Cx43 (left hand bar chart) and β1 (right hand bar chart) in untreated NRVMs and cells treated with βadp1 or scrambled βadp1 control peptide at 50 μM for 60 minutes n≥3 images per group. **F**) Representative confocal image of 1610 cells expressing β1 and GFP (green) immunolabeled for β1 (red). **G)** Diagram illustrating Electric Cell-Substrate Impedance Sensing (ECIS) and measurement parameters derived, including relative resistance of the monolayer overlying the electrode, which provides an assay of intercellular adhesion levels. **H)** ECIS comparison of effects on relativeresistance in monolayers of 1610 cells expressing β1, at 5 hours following treatment with vehicle control solution (n=6) or βadp1 or scrambled control βadp1 at 10 μM. *p<0.05, **p<0.01.

Confocal immunolabeling of NRVMs revealed β1 signals juxtaposed with Cx43 GJs at cell-to-cell contact sites in untreated cells (Fig. 1D) or cells treated with control peptide for 60 minutes consistent with localization at perinexal domains (Supplemental Fig. 1). This labeling occurred as sequential punctate domains of Cx43 and β1 signal at junctional contacts, which though intense and side-by-side, did not appear to directly co-localize or overlap. By contrast, following treatment of NRVMs with 50 μM βadp1 for 60 minutes, dissociation of juxtaposed side-by-side Cx43 and β1 signals was observed at junctional contacts (compare insets Fig 1D), together with qualitatively lower amounts of β1 immunolabeling. This reduction was confirmed by image quantification, which indicated that the density and counts of immunolabeled β1 at junctional contacts were significantly reduced in NRVMs exposed to βadp1 (Fig 1E). Junctional Cx43 immunolabeling density or counts did not show any significant change in response to βadp1 over 60 minutes, relative to controls. Similarly, and in line with whole cell sodium channel activity measurements, the total % areas of Cx43 and β1 immunolabeling normalized to cell area did not vary significantly between untreated cells, and NRVMS treated with 50 μM βadp1 or control peptide for 60 minutes (data not shown).

Further characterization of βadp1 over an acute treatment time-course was provided by electric cell substrate impedance sensing (ECIS) of 1610 cells stably transfected with SCN1B (β1) and GFP (1610β1 cells; Fig 1F). In ECIS, changes in relative resistance enable assay of levels of intercellular adhesion (Fig. 1G). Extending previous work, we confirmed that relative to vehicle and scrambled controls, a relatively low concentration of βadp1 (10 μM) was sufficient to prompt reductions in relative resistance/intercellular adhesion in 1610β1cells following 5 hours of exposure of the cells to the peptide (Fig 1H).

In sum, the results shown in Figure 1 indicate that disruption of β1-adhesion mediated by treatment with βadp1 over acute time courses of up to 5 hours is associated with reductions in junctional I_Na_ density and SCN1B (β1) immunolabeling levels, with global I_Na_ and SCN1B (β1) levels across the entire cell membrane remaining unaffected.

### Identification of Novel SCN1B Mimetic Inhibitory and Monomeric Agonist Peptides

Modeling *in silico* indicates that βadp1 likely mediates its effects over 30 to 60 minute time courses by selectively binding to the β1 extracellular Ig domain (Figure 2A, B). Experimental and modeling data further suggests that this interaction inhibits trans-adherent interactions between apposed β1 molecules on neighboring cell membranes [21, 39, 40]. Our approach to design of βadp1 as an inhibitor of β1-mediated adhesion was motivated in part by a strategy reported by groups working on two other cell adhesion molecules; N-cadherin [41] and desmosglein-2 [42]. These groups identified peptides in the Ig domain of these molecules that when provided in isolation inhibited adhesive interactions, but also described an approach to promoting adhesion based on dimerization of inhibitory monomers.

**Figure 2.**
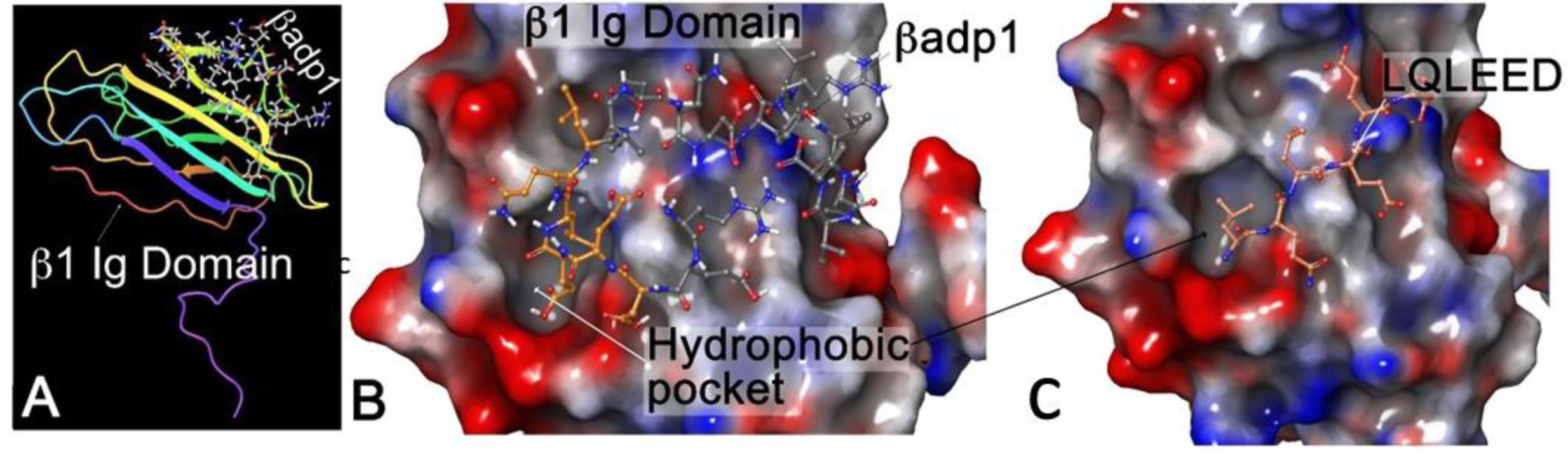
In silico modeling of binding of βadp1 and LQLEED monomeric inhibitory peptides to the β1 Immunoglobulin domain. **A)** Homology model of the β1 extracellular domain based on the SCN3B/β3 crystal structure [21], with βadp1 (stick model) docked *in silico* in a low-energy conformation with the adhesion surface of the β1 Immunoglobulin (Ig) loop. **B)** Rendering of βadp1 (ball and stick model) docked to thepredicted Ig adhesion surfaceof a β1 homology model. This perspective of the β1 Ig domain adhesion surface reveals a hydrophobic pocket, within which hydrophobic amino acids towards the C-terminus of βadp1 (e.g., LQL) are predicted to interact with β1. Positive and negatively charged moieties on the adhesion surface of the β1 Ig loop are shown in red and blue, respectively. **C)** Docking of CT sequence LQLEED from βadp1 with the β1 homology model in a low-energy conformation. The hydrophobic LQL sequence of peptide in this pose also embeds in the aforementioned hydrophobic pocket of the β1 Ig domain.

A second consideration of our approach to β1 drug design stemmed from data we obtained that molecules of greater than 3 kD do not efficiently penetrate the extracellular space of IDs in vivo [27]. The βadp1 monomer has a molecular mass of 2.6 kD and so does not exceed this threshold (Table 1). Thus, consistent with βadp1 having potential to target ID-localized β1 we found that the peptide had quantifiable effects on perinexus structure in Langendorf-perfused guinea pig hearts [21]. A dimeric peptide comprising βadp1 repeats exceeds the 3kD threshold, likely negating its ability to act as an ID-localized β1 agonist in the heart in vivo. As a first step in the generation of β1 agonist that could penetrate IDs, we used *in silico* modeling and ECIS to identify short peptides within βadp1 that had potential to interact with β1 and inhibit β1-mediated adhesion (Supplemental Fig. 2). Based on this approach we found that short peptides near the CT of βadp1 maintained propensity to interact with the β1 Ig domain and disrupt adhesion. Figure 2C shows one of these peptides, the 6mer LQLEED (Molecular mass ∼ 746 daltons), organized in a low energy pose with β1, with its NT leucine residues in a hydrophobic pocket on the Ig domain adhesion surface, within which portions of βadp1 also appear to embed (compare Figs. 2B and C).

**Table 1.**
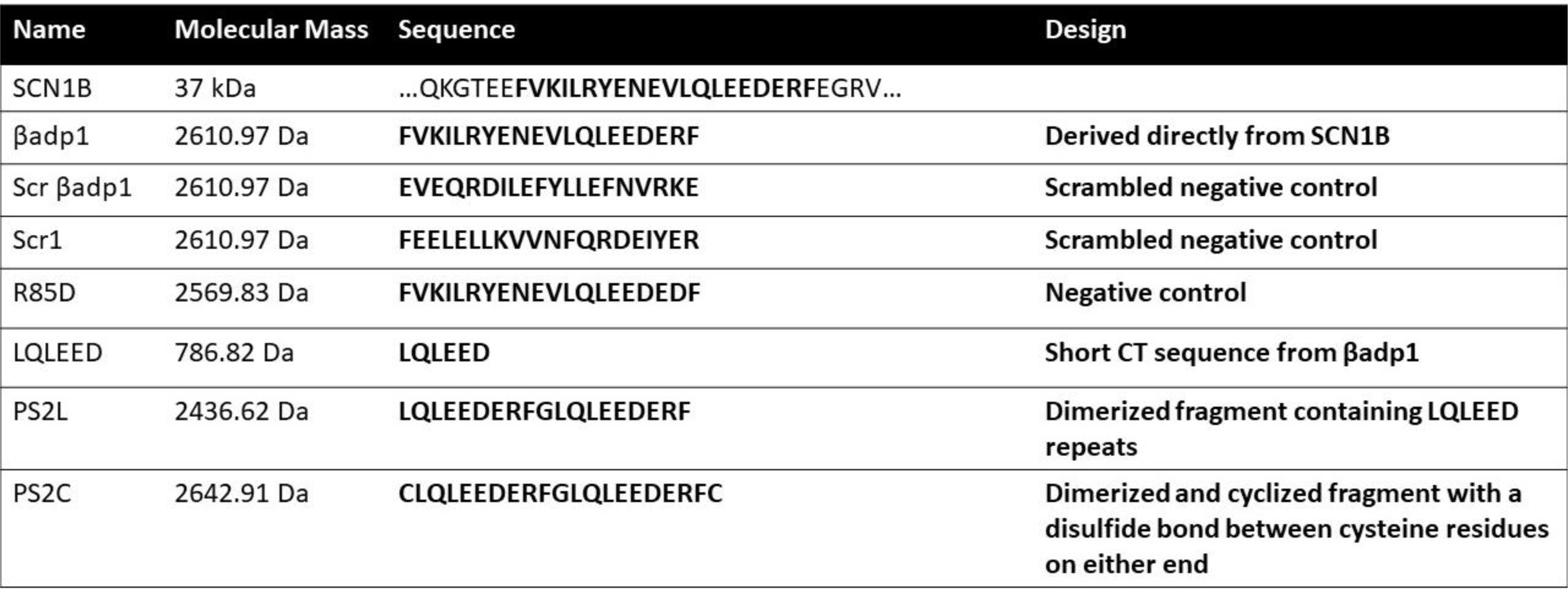

Next, we synthesized LQLEED, and applied the peptide at a 10 μM concentration in ECIS on 1610β1 cell monolayers (Figure 3). Similar to when low concentrations of βadp1 are used, we found that LQLEED caused a reduction in relative resistance like that observed for the larger 19 aa peptide – consistent with the peptide inhibiting intercellular adhesion between 1610β1 cells. By contrast, short peptides from nearer the βadp1 NT showed no evidence of similar inhibitory activity. (Supplemental Fig 2). The inhibitory effect of LQLEED is further illustrated in Figure 3A, where it is shown to reduce in relative resistance in 1610 monolayers following 5 hours of exposure of the cells to the peptide.

**Figure 3.**
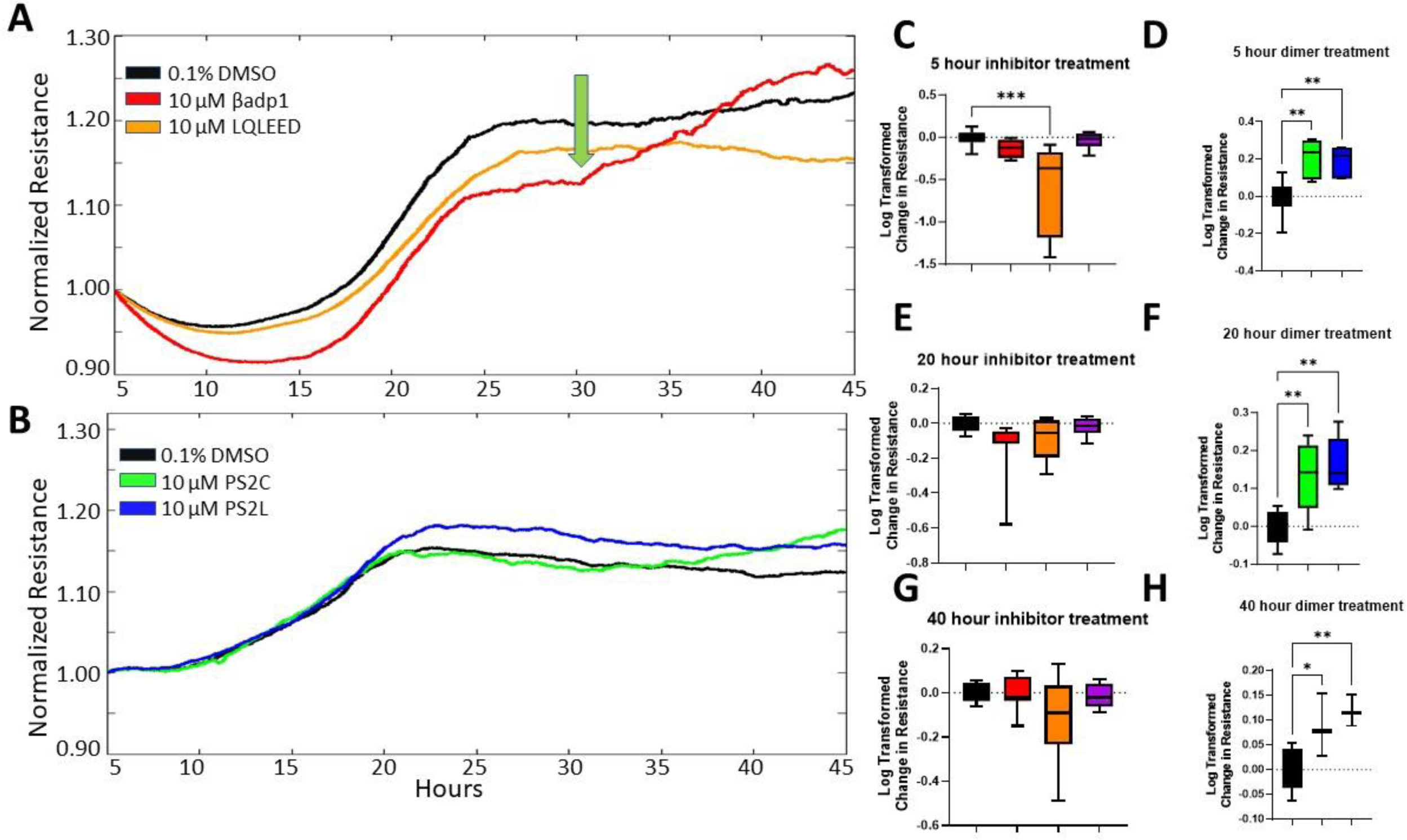
βadp1 decreases intercellular adhesion in 1610β1 cells, whereas βadp1-related dimeric peptides PS2C and PS2L increase adhesion. **A)** Multi-well ECIS showing effects of βadp1 (red) and the βadp1-derived peptide monomer LQLEED (orange) on intercellular adhesion as assayed by relative resistance in ECIS compared to vehicle (DMEM/F12 culture media with 0.1% DMSO) control treated (black) cells over 48 hours. Both βadp1 and LQLEED decrease adhesion compared to control at 5 and 20 hours of treatment. There is a shift in the curve of βadp1 at approximately 30 hours indicating increased adhesion is likely occurring beyond this timepoint (green arrow). As a result, at 40 hours there is no significant difference between βadp1 or LQLEED compared to the control. **B)** Multi-well ECIS demonstrating effects of βadp1-derived dimers, PS2C (green) and PS2L (blue) on intercellular adhesion compared to vehicle control treated (black) cells over 48 hours. Both PS2C and PS2L increase relative resistance/adhesion compared to controls at 5 and 20 hours. At 40 hours, PS2L continues to significantly increase adhesion. **C-H)** Quantification of ECIS relative resistance as compared to vehicle control following treatment with 10 μM βadp1 and 10 μM LQLEED inhibitory monomers at 5 **(C)**, 20 **(E)**, and 40 **(G)** hours after treatment or PS2C and PS2L agonizing dimers at 5 **(D)**, 20 **(F)**, and 40 **(H)** hours after treatment. n≥3 experimental replicates for each treatment, *p<.05, **p<.01.

To investigate whether dimerization of short CT inhibitory monomers could generate dimeric agonists, we synthesized LQLEEDERF-G-LQLEEDERF (Table 1). This dimeric peptide, which is called PS2L, is a sequential repeat of the CT-most 9 amino acids of βadp1 including the LQLEED sequence spaced by a glycine (G) linker. We also synthesized a dimeric peptide of identical sequence, except that it had cysteine residues at its NT and CT. This novel peptide, called PS2C (CLQLEEDERF-G-LQLEEDERFC), was designed to be cyclizable via disulfide bonding between its cysteines (Table 1). Importantly, the molecular mass of the PS2L and PS2C dimers are approximately 2395 and 2602 daltons, which like βadp1 and LQLEED falls below the 3 kD threshold for ID-penetration. Results from PS2L and PS2C will be described in detail in the following section – but to summarize in ECIS assays, resistance levels induced by these dimeric peptides in 1610β1 cells were notably elevated over controls (Figs. 3B).

### Differential Effects of Monomeric and Dimeric Mimetic Peptides Targeting β1 on Cell Adhesion and β1 Immunolabeling over Prolonged Time-courses

Whilst βadp1 and LQLEED reduced intercellular adhesion over courses of 5 hours or less, we found differential effects of the monomers on cellular adhesion over longer times periods. Up to 20-hours, the effects of monomers on cell adhesion were similar to that at 5 hours, with decreases in resistance, compared to controls (Figure 3A, C, E). However, after 24-30 hours a shift occurred, wherein resistance appeared to increase from the lower levels induced by the peptides acutely - as indicated by the green arrow in Figure 3A. In some ECIS runs, the rise in resistance in monomer-treated cells eventually exceeded that of control cells – as shown for βadp1 in Figure 3A. Whilst this increase did not always go above control levels, it was sufficient on average that by 40 hours post-treatment, effects of βadp1 and LQLEED on cellular resistance were no longer significantly below those of controls (Compare Figs. 3C and 3G). We performed similar long-term ECIS assays on the PS2L and PS2C dimeric peptides. Paradoxically with respect to the monomeric peptides, linear PS2L and cyclized PS2C dimers both significantly increased resistance in 1610β1 cells (Figs. 3D, F and H). The agonistic effects of the dimers were notably more potent than those elicited by the monomers, reaching both higher levels of significance and persisting over the entire 40 hour time course of the experiment.

Immunolabeling was undertaken to probe the effect of the mimetic peptides on 1610β1 cells over a 48 hour time course using an antibody to the NT Ig domain of β1 (Fig. 4). 1610β1 cells incubated with 50μM Biotin-βadp1 showed accumulation of the peptide, which peaked at around 24 hours, declining at 48 hours. Interestingly, immunolabeling for the β1 protein itself indicated a steady increase in abundance in 1610β1 cells in response to βadp1, which reached maximal levels at 48 hours, relative to control cells (Fig. 4). Prompted by this observation we examined what effects that LQLEED and PS2L had on β1 immunolabeling at 48 hours (Fig. 5A-E). As was the case for βadp1, in response to LQLEED β1 immunolabeling in 1610β1 cells appeared to be increased at 48 hours (Fig. 5C). High magnification images indicated that this increased immunolabeling was cell-wide, including within the cytoplasm, with occasional evidence of increased intensity of immunolabeling at cell borders. PS2L treatment also led to cell-wide increases in β1 immunolabeling at 48-hours. Interestingly, we also noted that treatment with PS2L prompted a distinct increase in signal at cell borders, relative to controls and the monomeric peptides (Fig. 5D, E). The increase in cell border localization observed with the dimeric peptide was not seen for the monomers.

**Figure 4.**
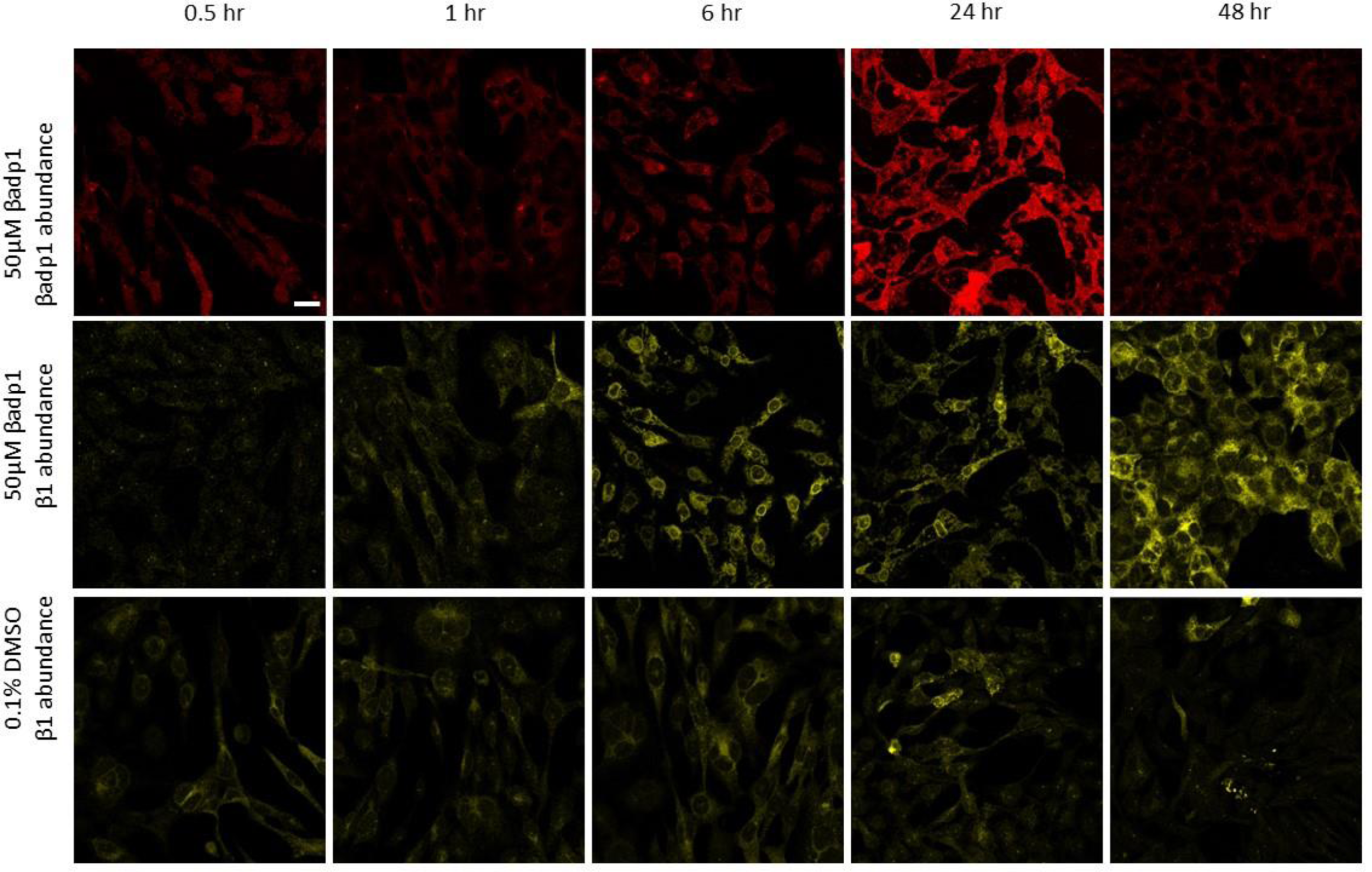
βadp1 increases β1 immunolabeling over a 48 hour Time-Course. Representative images of 1610β1 cultures treated with 10 μM biotinylated βadp1 or vehicle control over 48 hours, with sampling at 0.5, 1, 6, 24 and 48 hours. The top two rows are paired double-labelings of fluor-labeled streptavidin and β1 immunolabeling within the same image. Biotin-βadp1 levels peak at 24 hours during the time course, with prominent intracellular signal in evidence. β1 immunolabeling abundance in the 1610β1 cells appears to peak 24 hours later at the 48 hour time point. The bottom row indicates that β1 immunolabeling abundance remains relative steady over time course in control cells.

**Figure 5.**
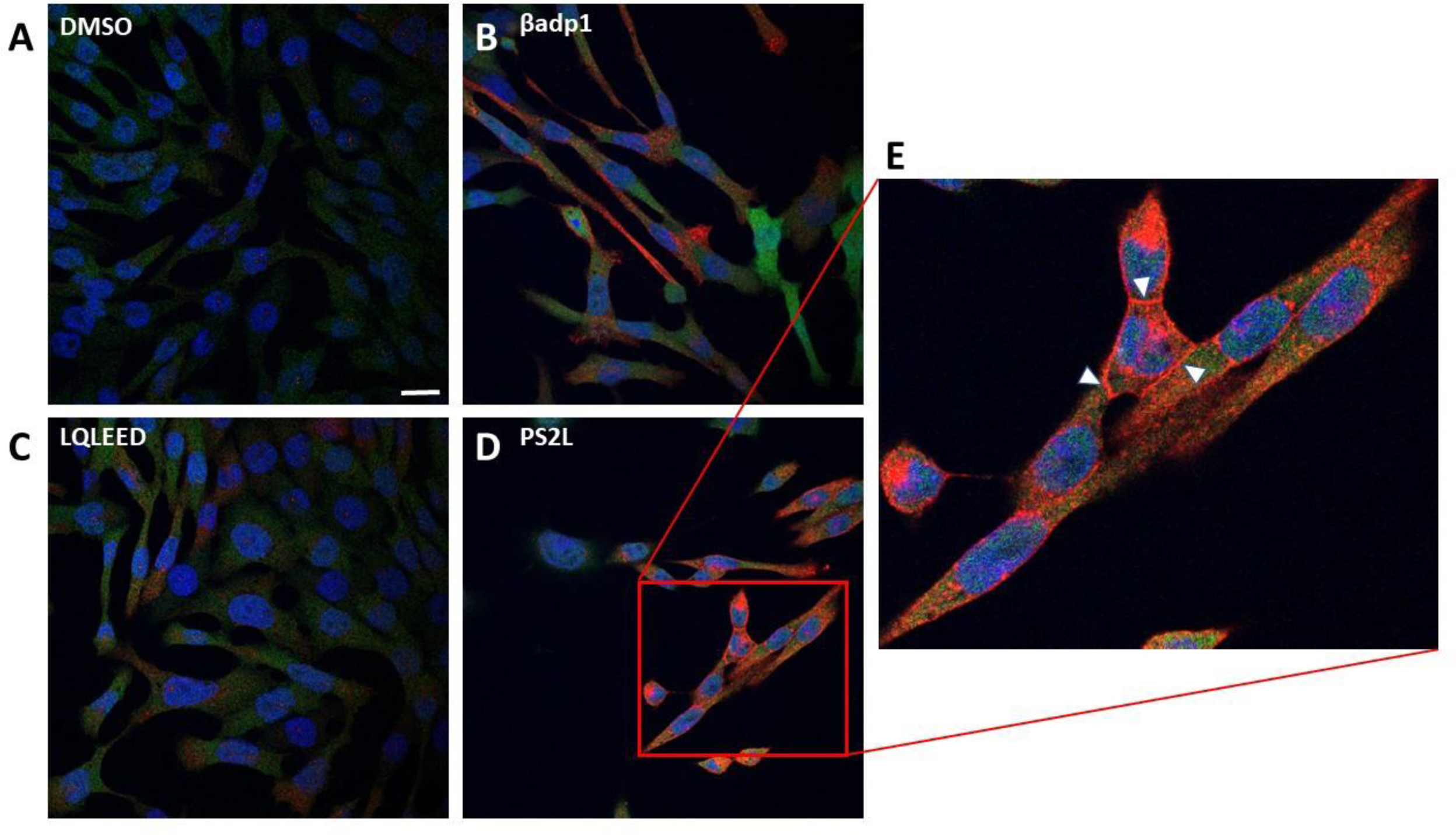
βadp1 and βadp1-derived monomeric and dimeric peptides increase β1 immunolabeling at 48 hours of treatment. 1610β1 cells treated for 48 hours with vehicle control solution **(A)** or 10μM βadp1 **(B)** or the βadp1-derived peptides LQLEED at 10μM **(C)** or PS2L at 10μM **(D).** Each peptide shows an increase in β1 immunolabeling abundance (red) compared to vehicle control cells at 48 hours. **E)** Magnified view of indicated portion of PS2L-treated cells. PS2L increased overall β1 abundance in 1610β1 cells, but unlike the other βadp1-derived peptides, it also increased the β1 abundance at cell-cell borders (white arrows). Scale bar: 10μM.

### βadp1 and PS2L Increase Cleavage of β1 via Regulated Intramembrane Proteolysis (RIP)

β1 has been reported to undergo a process of sequential intramembrane proteolysis (RIP) by BACE1 and γ-secretase [36, 38]. RIP results in the production of a soluble intracellular domain (ICD) from the CT of full length β1 that is translocated to the nucleus, with correlated effects on gene expression, including VGSC sub-units. We sought to determine if β1-targeting peptides effect the RIP process in a manner that may parallel changes in β1-mediated intercellular adhesion and immunolabeling. First, to confirm that β1 underwent RIP in 1610β1 cells, we treated cells with the γ-secretase inhibitor DAPT. In the presence of DAPT, together with an antibody against the CT of β1, it was found that the 19 kD Carboxyl-Terminal Fragment (CTF) accumulated in Western blots from cells sampled at 6, 24 and 48 hours following initiation of treatment (Fig. 6A). Without DAPT treatment the 19 kD CTF was typically undetectable. Next, we co-treated cells with DAPT in the presence of βadp1 (Fig. 6). As expected, all treatments that included DAPT resulted in increased levels of the CTF (Fig. 6A). However, compared to DAPT alone, βadp1+DAPT significantly increased CTF levels at 6, 24, and 48 hours post treatment (Fig. 6B). Cells treated with a control peptide showed no similar increase in the 19 kD CTF (Supplemental Fig 3). We noted that βadp1 alone increased the 19 kD fragment at 24 hours of treatment relative to control levels on occasion (Fig. 6A). However, this was not consistent between experiments and we did not analyze this sporadic effect further. The abundance of the full length β1 (37 kD band) was elevated at 6 hours post-treatment by βadp1 in DAPT co-treatments (Fig. 6C), but other than this modest effect, the 37 kD band showed no consistent variation by treatment or time-point on Western blots.

**Figure 6.**
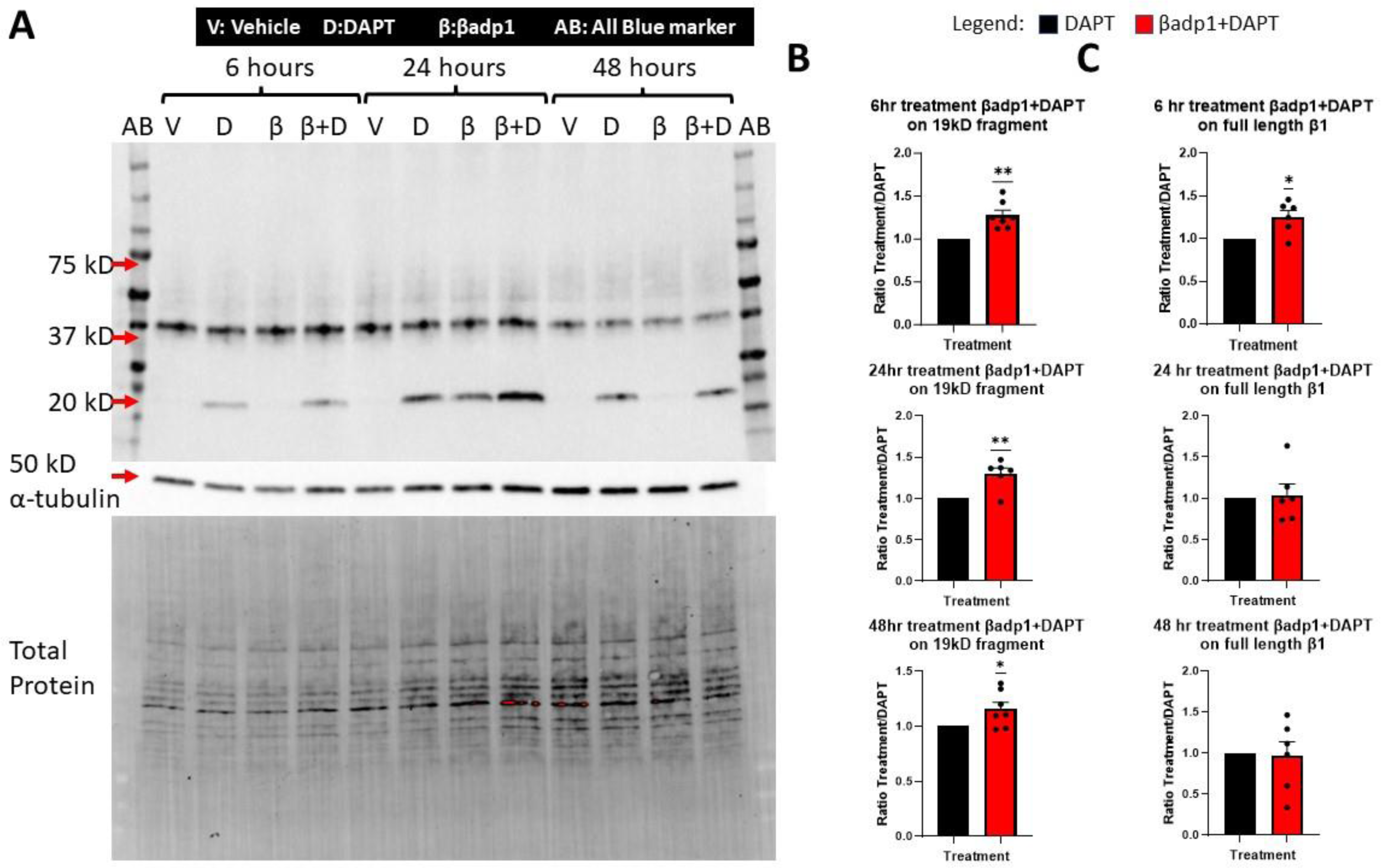
βadp1 increases Regulated Intramembrane Proteolysis (RIP) of β1 following 6, 24 and 48 hours of treatment. **A-C)** Western blot analysis demonstrating βadp1 effects on β1 and the 19 kD β1 Carboxyl Terminal Fragment (CTF) generated by RIP. Blot shows effects following treatment with vehicle control (DMEM/F12 culture media with 0.1% DMSO), 1 μM DAPT, 50μM βadp1, 50μM βadp1 +1μM DAPT at 6, 24 and 48 hours of treatment. Below the β1 immunoblot in **(A)** are α-tubulin and total protein loading controls, showing approximately equal loading in each lane. **B)** Quantification of the β1 CTF at 6, 24, and 48 hours normalized to vehicle control, indicating significant increases in the 19kD fragment when βadp1 treatment is present together with DAPT, relative to DAPT alone at the 6, 24 and 48 hours of a two-day treatment time course. DAPT inhibits γ-secretase, leading to cytoplasmic accumulation of the 19 kD β1 CTF, the product of BACE-1-mediated proteolysis, enabling resolution of βadp1 effects on RIP. **C)** Quantification of full-length β1 normalized to vehicle control in the presence of DAPT or βadp1 plus DAPT over the same time course. There is a significant increase in full-length β1 6 hours after co-treatment with βadp1 and DAPT, but no significant difference in levels of β1 persist at 24- and 48-hours. n≥3 experimental replicates for each treatment, *p<.05, **p<.01.

PS2L+DAPT also significantly increased the CTF of β1 at 6 hours post treatment compared to DAPT alone, though not at 24 and 48 hours (Fig. 7). PS2L alone did not appear to have a consistent effect on CTF levels. In response to combinatorial DAPT and PS2L treatments, full length β1 showed no change at the 6 and 24 hour time points, though at 48 hours Western blots revealed a significant reduction of the 37 kD band relative to controls.

**Figure 7.**
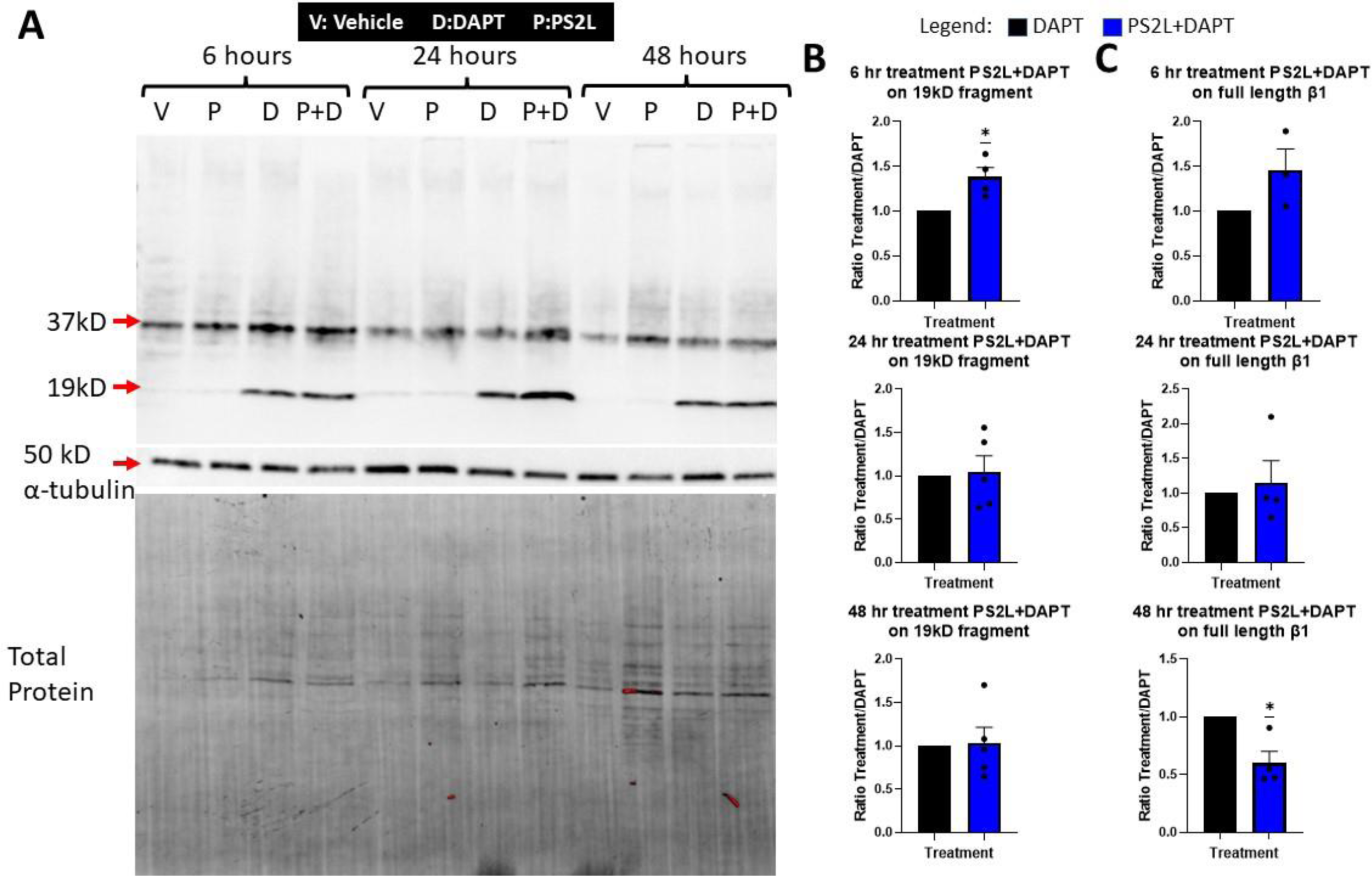
The PS2L dimer acutely increases the RIP of β1, but not subsequently. **A-C)** Western blot analysis showing effects on full-length β1 and the β1 CTF of treatment with vehicle control (DMEM/F12 with 0.1% DMSO), 1 μM DAPT, 50μM PS2L, 50μM PS2L +1μM DAPT for 6, 24 and 48 hours. Below the β1 immunoblots in **(A)** are a-tubulin and total protein controls, showing approximately equal loading in each lane. **B)** Quantification of the β1 CTF at 6, 24, and 48 hours normalized to vehicle control, indicates a significant increase in the 19kD fragment occurs at the 6 hour time-point of a two-day treatment time course in response PS2L treatment in the presence of DAPT, but not at 24 and **C)** Quantification of full-length β1 normalized to vehicle control in the presence of DAPT or PS2L plus DAPT over the same time course. There is a significant decrease in full-length β1 following 48 hours of co-treatment with βadp1 and DAPT. n≥3 experimental replicates for each treatment, *p<.05, **p<.01.

To explore whether inhibiting β1 RIP may disrupt the longer term gain-of-function effect of βadp1 on intercellular adhesion in 1610β1 cells, we repeated ECIS experiments with βadp1 treatment over 48 hours, including a βadp1 + DAPT treatment (Fig. 8C). We found that a co-treatment of βadp1 and DAPT resulted in prolongation of the inhibitory effect, with relative resistance significantly decreased across the entire length of the experiment (Fig. 8D-F). These results would be consistent with contribution of RIP to the longer term gain-of-function effect on intercellular adhesion observed with treatment with βadp1 over periods of greater than 24 hours.

**Figure 8.**
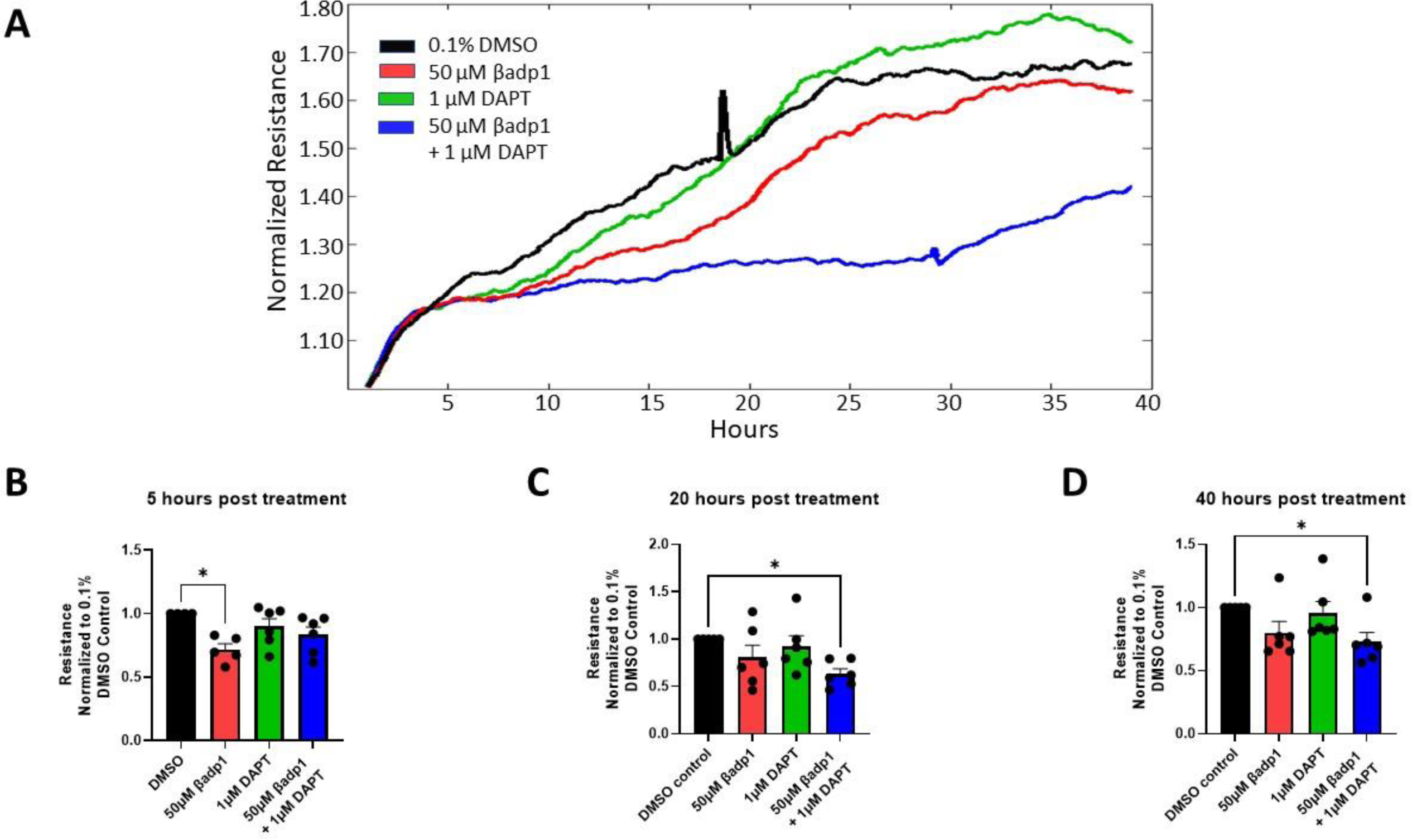
DAPT inhibition of RIP results in loss of βadp1 effects on ECIS-assayed cell adhesion. **A)** Multi-well ECIS demonstrating activity of βadp1 (red), DAPT (green), and βadp1+DAPT (blue) compared to vehicle control (black) treated 1610β1 cells. **B-D)** Quantification of ECIS data at 5-, 20-, and 40-hours post-treatment. βadp1 repeats previous results, showing a decrease in resistance at 5 hours, but then steadily increases up to the 40-hour timepoint with no significant difference between control and βadp1 resistance at 20 and 40 hours. DAPT does not significantly alter the resistance at any timepoint. However, βadp1+DAPT decreases resistance compared to DMSO across the entire experiment, suggesting that inhibition of the RIP of β1 disrupts the longer-term gain of adhesion mediated by βadp1 beyond 24 hours.

**Figure 9.**
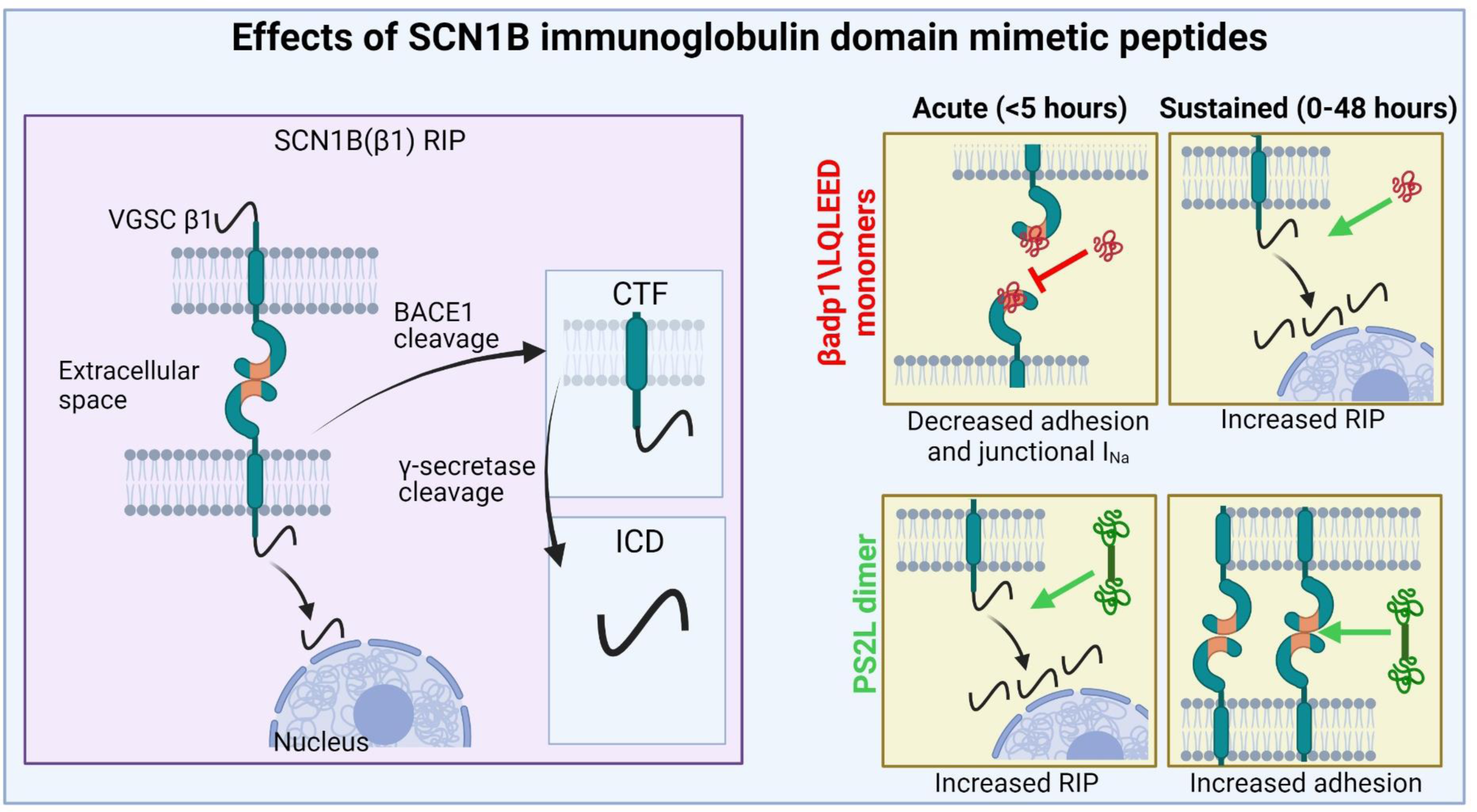
Model for Effects of SCN1B (β1/β1B) immunoglobulin domain mimetic peptides. The left panel illustrates the adhesion function and regulated intramembrane proteolysis (RIP) of SCN1B (β1). Full length β1 is 37kD and is first cleaved by BACE1. This releases the extracellular domain of β1 containing the Ig domain responsible for cell adhesion, resulting in a 19kD fragment simply called the c-terminal fragment (CTF), which is still located in the membrane and is made up of the transmembrane and intracellular portions of β1. The CTF is sequentially cleaved by γ-secretase. This results in the intracellular domain (ICD) being released. The ICD is the final cleavage product and translocates to the nucleus, resulting in transcriptional changes. The 4 yellow panels to the right illustrate the effects of the βadp1/LQLEED monomers and the PS2L dimer on adhesion and RIP during acute and sustained treatments. βadp1/LQLEED decrease adhesion and junctional I_Na_ acutely while increasing RIP over sustained treatment. PS2L increases RIP on an acute level and increases adhesion over the sustained treatment timecoures. Figure created with BioRender.com.

## DISCUSSION

In the present study, the effects of targeting VGSC β1 with amino acid sequences mimicking its extracellular immunoglobulin (Ig) domain were investigated. A short sequence (LQLEED) was characterized at the C-terminus of βadp1 - a previously reported 19 amino acid mimetic of the SCN1B Ig domain [21]. LQLEED inhibited cell-to-cell adhesion in 1610 β1-expressing cells in a manner comparable to βadp1. Dimeric peptides incorporating repeats of LQLEED were found to demonstrate agonist-like activity, paradoxically increasing adhesion in 1610 β1 monolayers in contrast to the inhibitory effects of monomeric peptides. New insights into the activity of SCN1B mimetic peptides over time courses longer than previously reported were provided, including that: 1) The βadp1 monomer reduced peak sodium current at intercellular contacts between cardiomyocytes, associated with reductions in β1/β1B immunolabeling at these junctional regions; 2) The inhibitory effects of monomeric βadp1 and LQLEED on cell adhesion in 1610 β1 expressing monolayers was biphasic, being sustained for time courses of up to 5 hours, but thereafter blunted; 3) Conversely, the adhesion-promoting effects of dimeric peptides were maintained over 48 hours; 4) The biphasic time course of βadp1 on cell adhesion in 1610 β1-expressing cells was correlated with increased levels of a C-terminal fragment (CTF) of β1 – a known intermediate product of the regulated intramembrane proteolysis (RIP) of β1 [36, 38] and that 5) The second phase of the biphasic effect of βadp1 appeared to be prevented by co-treatment with DAPT - an inhibitor of γ-secretase a peptidase involved in the final step of β1 RIP [36, 38]. In sum, our results suggest pathways to pharmacological development of agonists and antagonists targeting the main beta subunit (i.e., SCN1B) of the principle sodium channel found in the mammalian ventricle [13, 17], as well as complexities in the response of β1-expressing cells to such pro-drugs.

When designing the peptides derived from the initial βadp1 sequence, we ensured biologically relevant residues were included. A human population study of SCN1B mutants found an R85H variant in a patient with atrial fibrillation [34]. When the mutant β1 subunit was expressed in Chinese hamster ovary cells, sodium current via Na_v_1.5 was reduced [34]. Interestingly, this mutation is also associated with epilepsy [14, 43]. The full βadp1 sequence is derived from residues 67-86 of β1, which includes amino acid R85.

Another mutation one residue removed from the βadp1 sequence, E87Q, has been associated with Brugada syndrome [35]. Thus, combined with our *in silico* modeling and experimental ECIS results (Supplemental Fig. 2) we focused on the carboxyl-terminus (CT) of βadp1 when developing short dimeric and monomeric mimetic sequences of the β1/β1B Ig domain. Interestingly, the control peptide βadp1-R85D, does not appear to cause significant changes in accumulation of the CTF compared to DMSO, unlike βadp1 – again reinforcing the concept that bioactivity resides in the CT region of the sequence (Supplemental Fig. 3).

Although the focus is on β1 in this study, there is an alternatively spliced variant of SCN1B named β1B. β1B is formed by retention of an intron that results in a novel CT that lacks the transmembrane domain of β1 [44, 45]. Thus, β1B is not thought to be anchored in the cell membrane and is widely held to be secreted, except in one report where β1B was found to be retained in the cell membrane when co-expressed with Na_v_1.5 in HEK cells [14]. Since β1B is exclusively retained at the membrane in the presence of Nav1.5, the predominant cardiac VGSC isoform, this suggests a unique role for β1B in the heart, including potential to act as an adhesive substrate. Although β1B differs at the CT from β1, it does contain the same Ig domain that appears to be necessary for adhesive interactions mediated by the canonical (i.e. the non-alternatively transcribed) β1 isoform. This leads to the question of whether βadp1 could mimic a natural function of β1B in regulating β1-mediated cell adhesion, as well as modulating β1 RIP, since βadp1 is a sequence common to both β1B and β1. Interestingly, β1B loss of function mutations are implicated in various cardiac and neural pathologies [46], including Long QT-Syndrome [47], epilepsy [48, 49], Dravet syndromes [50, 51], Brugada syndrome [35, 52], and atrial fibrillation [34, 53]. Importantly, the R85D and E87Q mutations near or within the βadp1 sequence can occur in both β1 and β1B. Further work is needed to determine the differing roles of β1 and β1B and in regard to the SCN1B (β1/β1B) peptides studied herein.

To further understand how short peptides mimicking the β1/β1B Ig domain prompt changes to β1-mediated adhesion, we investigated effects of our peptides on RIP of β1. β1 RIP occurs in two steps. First, BACE1 cleaves the extracellular Ig domain, leaving the 19kD CTF composed of the transmembrane and intracellular domains [36, 38]. Under normal conditions, the 19kD CTF is subsequently cleaved by γ-secretase to produce an intracellular domain (ICD) that translocates to the nucleus and results in transcriptional changes, including differential changes in gene expression of VGSC genes, as well as genes responsible for cell adhesion, immune response, cellular proliferation, and calcium ion binding [38]. It has been shown in MDA-MB-231 cells, a breast cancer cell line, that over-expression of full length β1-GFP or ICD-GFP alone is sufficient to increase sodium currents, whereas expression of β1STOP-GFP, which does not include the ICD, does not increase sodium current [54]. This indicates that the SCN1B mimetic peptides that are the focus of the present study may be capable of modulating transcriptional effects mediated by the ICD, as well as causing changes to sodium currents. Whether or not peptides such as βadp1 and LQLEED have this potential represents a fruitful line of inquiry for ongoing study.

Aside from β1, there are multiple other substrates of similar RIP processes, including the β-amyloid precursor protein (APP) implicated in Alzheimer’s disease and Notch. The cleavage process of APP that produces Aβ plaques involves sequential activity of β-secretase and γ-secretase. The resulting intracellular domain (AICD) also shows nuclear signaling activity, including effects such as endogenous defense against Alzheimer’s disease [55] and regulating neurogenesis [56]. Notch is a transmembrane protein that is sequentially cleaved by ADAM10 then γ-secretase once it is in the membrane [57]. Comprehending the much larger body of work surrounding RIP of both APP and Notch, as well as other voltage gated ion channel subunits and cell adhesion molecules, appears to offer the prospect of further progress on understanding the β1 RIP process and targeting it for therapeutic purposes [58].

One potential downstream method by which increasing β1 RIP results in increases in cellular adhesion as seen in our various ECIS assays, may be via increased abundance of the β1 subunit itself. The literature has not shown whether the β1 ICD affects levels of VGSC β subunit transcription, but it does upregulate VGSC α subunit transcription [36, 38]. Localization of α and β subunits are closely related, with β subunits usually found in regions of high α subunit density [59]. Our immunofluorescent data also indicates an increase in β1 abundance in the 1610β1 cells after treatment with SCN1B (β1/β1B) mimetic peptides. Previously, we have shown that β1 is important in maintaining inter-membrane adhesion at the perinexus, a specialized nanodomain directly adjacent to gap junctions in the intercalated disc that is the proposed region where ephaptic coupling takes place in cardiac tissues [21]. Knockout of the β1 protein in mice is fatal by postnatal day 21 [60] and disrupting β1 trans-adhesion in the perinexus *ex vivo* results in loss of GJ-associated VGSCs, conduction slowing and arrhythmias [21]. By instead increasing the levels of β1, these rationally designed peptides may thus have antiarrhythmic treatment or prevention abilities, albeit that our present data suggests that there may be acute pro-arrhythmic effects of such approach to therapy. Ongoing studies may well address the latter issue, but until such time it is probably best that assessments of the potential long-term clinical benefits of such strategies be taken with caution.

One question raised by our studies, and by others, is whether or not RIP of β1 takes place at the cell membrane or subcellularly. Our immunofluorescent data indicates that β1 heterologously expressed in 1610 cells is mainly cytoplasmic. This observation is similar to those made by others in the MDA-MB-231 β1 expressing cells [54] and Madin-Darby canine kidney cells [61]. In the MDA-MB-231 cells β1 has been shown to colocalize with the endoplasmic reticulum, the endolysosomal pathway and the nucleus [54]. From our immunofluorescence experiments we find that there is a significant uptake of βadp1 into the cells at 24 hours post-treatment. The greatest increase in β1 abundance occurs at 48 hours post-treatment, which aligns with the increase we see in resistance in ECIS. There is little difference in β1 levels between DMSO and βadp1-treated cells up to this point. We also see the greatest increase of CTF with βapd1+DAPT at 24 hours post-treatment, corresponding to the timing of maximal uptake of the peptide by cells. This suggests that the internalization of the peptide is important in causing the observed increase in RIP. Thus, the literature, together with data presented here, suggests that a considerable fraction of β1 RIP may occur intracellularly, so βadp1 may affect the process to a greater extent once it has been taken up by the cells. This indicates potential for delivery systems such as small extracellular vesicles loaded with SCN1B (β1/β1B) mimetic peptides that can be directly delivered inside the cells. Previous work has shown potential for drug delivery with bovine milk derived extracellular vesicles, and our lab has optimized a protocol for isolating large quantities of small extracellular vesicles to allow for loading and treatment of the vesicles [62–64]. This being said, evidence remains that the majority of β1 RIP can and does occur at the plasma membrane [37, 54]. β1 is S-palmitoylated, and when palmitoylation is inhibited by a C162A mutation in β1, this results in decreased β1 in the plasma membrane and decreased RIP [37]. Therefore, ongoing work is needed to establish the location of β1 RIP in different biological settings, both in health and disease and to compare efficacy of extracellular versus intracellular delivery systems for peptides targeting SCN1B.

We have previously hypothesized the function of β1 in the perinexus is to regulate intermembrane spacing via trans-adhesion and our peptides were initially designed with perinexal trans-interaction in mind [21]. While the initial and late effects of the βadp1 monomer may be respectively explained by an acute disruption of trans-adhesion among β subunits, and possibly a subsequent increase in SCN1B level via downstream effects on RIP, the basis for the sustained increase in resistance (and thus intercellular adhesion) in ECIS after treatment with dimeric PS2L awaits mechanistic explanation. Other groups have demonstrated that dimer mimetics of adhesion molecules result in increased adhesion, namely in desmoglein-2 and N-cadherin [41, 42]. However, if PS2L increased adhesion by facilitating trans-adhesion among β1 subunits, this would be at odds with how the increased RIP of β1 is triggered by both βadp1 and PS2L, with one inhibiting and one facilitating trans-adhesion. Therefore, more work is required to determine how PS2L facilitates adhesion while simultaneously increasing the rate of β1 RIP at early timepoints.

In summary, we show that β1 mimetic peptides increase the amount of regulated intramembrane proteolysis that β1 undergoes in 1610β1 cells. Although we do not provide evidence that the β1 mimetics studied herein directly affect transcription of β subunits of VGSCs, we see evidence that β1 abundance is increased in treated cells and previous work has shown changes to VGSC α subunit transcription in response to β1 ICD increase [38]. We also as yet have no direct evidence that increased resistance in ECIS corresponds to decreased perinexal width and changes to heart rhythm, so these are next steps. To conclude, we have identified novel SCN1B (β1/β1B) mimetic peptides with potential to inhibit or promote intercellular β1-mediated adhesion, possibly including by effects on β1 RIP, suggesting paths to development of anti-arrhythmic drugs targeting the perinexus.

## MATERIALS AND METHODS

### Peptide synthesis

Peptides used in the study are summarized in Table 1. Peptides were synthesized by and obtained from LifeTein (Somerset, NJ) modified by N-terminal acetylation, C-terminal amidation, and TFA removal. Peptides were solubilized in DMSO to the concentration shown by experiment, ranging from 10μM-100μM.

### Neonatal rat ventricular myocyte (NRVM) isolation

NRVM isolation was performed as previously described [21]. NRVM isolation procedures conformed to the UK Animal Scientific Procedures Act 1986. In brief, isolated ventricles were taken from one-day-old rat pups anesthetized with a lethal dose of isoflurane. The ventricles were sectioned into small cubes and processed using mechanical dissociation (gentleMACS) and enzymatic degradation (neonatal heart dissection kit; Miltenyi Biotec, Bergisch Gladbach, Germany). The cell suspension was then filtered and the cells were plated onto glass-bottom dishes (MatTek Corp., Ashland, MA) in M199 media supplemented with newborn calf serum (10%), vitamin B12, glutamate and penicillin/streptomycin (1%). Myocytes were allowed to grow and establish connections for 3–4 days *in vitro*. Peptide treatments were applied to cell monolayers the specified amount of time (Fig. 1) prior to I_Na_ measurements.

### Scanning ion conductance microscopy (SICM) guided smart patch clamp

A type of SICM, called hopping probe ion conductance microscopy, was combined with cell-attached recordings of cardiac sodium channels from NRVM monolayers, as previously described [21], to assess I_Na_ at cell-to-cell contact sites. Currents were recorded in cell-attached mode using an Axopatch 200A/B patch-clamp amplifier (Molecular Devices, Sunnyvale, CA), and digitized using a Digidata 1200B data acquisition system and pClamp 10 software (Axon Instruments; Molecular Devices, Sunnyvale, CA). Cell-to-cell junctional sites were identified using a sharp scanning nano-probe (40–50 MΩ), followed by controlled increase of pipette diameter (20–25 MΩ) for capture of active sodium channel clusters. The external solution contained (in mM): KCl 145, Glucose 10, HEPES 10, EGTA 2, MgCl_2_ 1, and, CaCl_2_1 (300 mOsm and pH 7.4). The internal (pipette) solution contained (in mM) NaCl 135, TEA-Cl 20, CsCl 10, 4AP 10, Glucose 5.5, KCl 5.4, HEPES 5, MgCl_2_ 1, CaCl_2_ 1, NaH_2_PO_4_ 0.4 and CdCl_2_ 0.2 (pH 7.4). The voltage-clamp protocol was made up of sweeps testing potentials from −70 to +30 mV from a holding potential of −120 mV.

### Molecular modeling

Molecular modeling was used as described previously to simulate the docking of βadp1 and LQLEED with the homology model of β1 using Maestro (Maestro, Schrödinger, LLC, New York, NY) [21].

### Whole-cell I_Na_ recordings in NRVMs

Whole cell sodium current recordings were collected as previously described [21]. Whole-cell I_Na_ was recorded from NRVMs in low-sodium extracellular solution containing the following (in mM): CsCl 130, NaCl 11, Glucose 10, HEPES 10, MgCl_2_ 2, CaCl_2_ 0.5, CdCl_2_ 0.3, adjusted to 7.4 with CsOH. The intracellular (pipette) solution contained (in mM): Cesium methanesulfonate (CsMeS) 100, CsCl 40, HEPES 10, EGTA 5, MgATP 5, MgCl_2_ 0.75, adjusted to pH 7.3 with CsOH. Pipettes were pulled from the borosilicate glass microelectrodes and had a resistance of 3–4 MΩ. Sweeps were initiated from the holding potential of −100 mV to test potentials ranging from −75 to + 20 mV in 5 mV increments. Peak current was measured between −30 and −40 mV. Whole cell capacitance ranged between 9 and 16 pF. Current density (in pA/pF) was calculated as the ratio of the peak current to cell capacitance.

### Heterologous expression of β1 in 1610 cells

We have previously described how Chinese hamster lung 1610 cells have been used as a heterologous expression system [21]. These cells were used to measure SCN1B/β1 mediated adhesion and the regulated intramembrane proteolysis of SCN1B/β1 and they do not endogenously express SCN1B/β1 [65].

### Electric cell-substrate impedance sensing (ECIS)

Resistance of adherent monolayers of 1610 cells expressing SCN1B/β1 was measured using an ECIS Zϴ system (Applied Biophysics) over the entire range possible, (62.5-64000 Hz). For analysis, we used the resistance calculated from the 4000 Hz measurement as previously shown [21]. Cells were plated on 8-well dishes (8W10E+, 40 electrodes per well, Applied Biophysics) at a concentration of 500,000cells/mL, 300μL per well. Impedance was measured continuously over the treatment time courses, and resistance is calculated as the real component of impedance.

### Fluorescent Immunolabeling

Fluorescent immunolabeling was performed on adherent monolayers of 1610 cells expressing SCN1B/β1 fixed with 4% paraformaldehyde for 10 minutes. Samples were labeled with a rabbit polyclonal antibody against an amino-terminal region of β1 (Epitope: _44_KRRSETTAETFTEWTFR_60_). We have previously published validation results for this antibody [21]. Samples were then labeled with goat anti-rabbit AlexaFluor 568 (1:2000, ThermoFisher Scientific) secondary antibodies for confocal microscopy. Nuclei were stained with Hoescht 33342 for 10 minutes (1:30,000, Invitrogen).

### Confocal microscopy

Confocal microscopy was done with a TCS SP8 laser scanning confocal microscope equipped with a Plan Apochromat 63x/1.4 numerical aperture oil immersion objective and a Leica HyD hybrid detector (Leica) using approach reported previously [21]. Imaging of each fluorophore was performed sequentially, and the excitation wavelength was switched at the end of each frame.

### Western blotting

Whole cell lysates were collected from 1610 cells expressing SCN1B/β1 using RIPA buffer on ice while being agitated on a platform rocker. Lysates were then pulled through 22 gauge and 17 gauge syringes 5 times each, vortexed, and spun for 30 minutes at 10,000g at 4°C before being snap-frozen in liquid nitrogen. Lysates were electrophoresed on 4-20% TGX Stain-free gels (BioRad) and were then transferred to polyvinylidene difluoride (PVDF) membrane using a semi-dry method with a Trans-Blot Turbo system for 7 minutes at 25V (BioRad). The membranes were probed with the rabbit polyclonal antibody against an amino-terminal region of β1 [21], or a rabbit polyclonal antibody against the carboxy-terminal of β1 (Cell Signaling Technology, D4Z2N) each diluted at 1:1000. These were followed by a goat anti-rabbit HRP-conjugated secondary antibody (JacksonImmuno). Signals were then detected using SuperSignal West Femto Maximum Sensitivity Substrate (ThermoFisher Scientific) and imaged with a ChemiDoc MP imager (BioRad).

### Statistical analysis

Data is presented as mean ± SEM unless specified. Statistical analysis was performed using one-way ANOVA with Dunnett’s multiple comparisons test or one-sample students t-test with a theoretical mean of 1 in the case of the Western Blot data. All statistical analysis was performed using GraphPad Prism v.10.0.2. A p<0.05 was considered statistically significant.

## SUPPLEMENTAL FIGURE LEGENDS

**Supplemental Figure 1.**
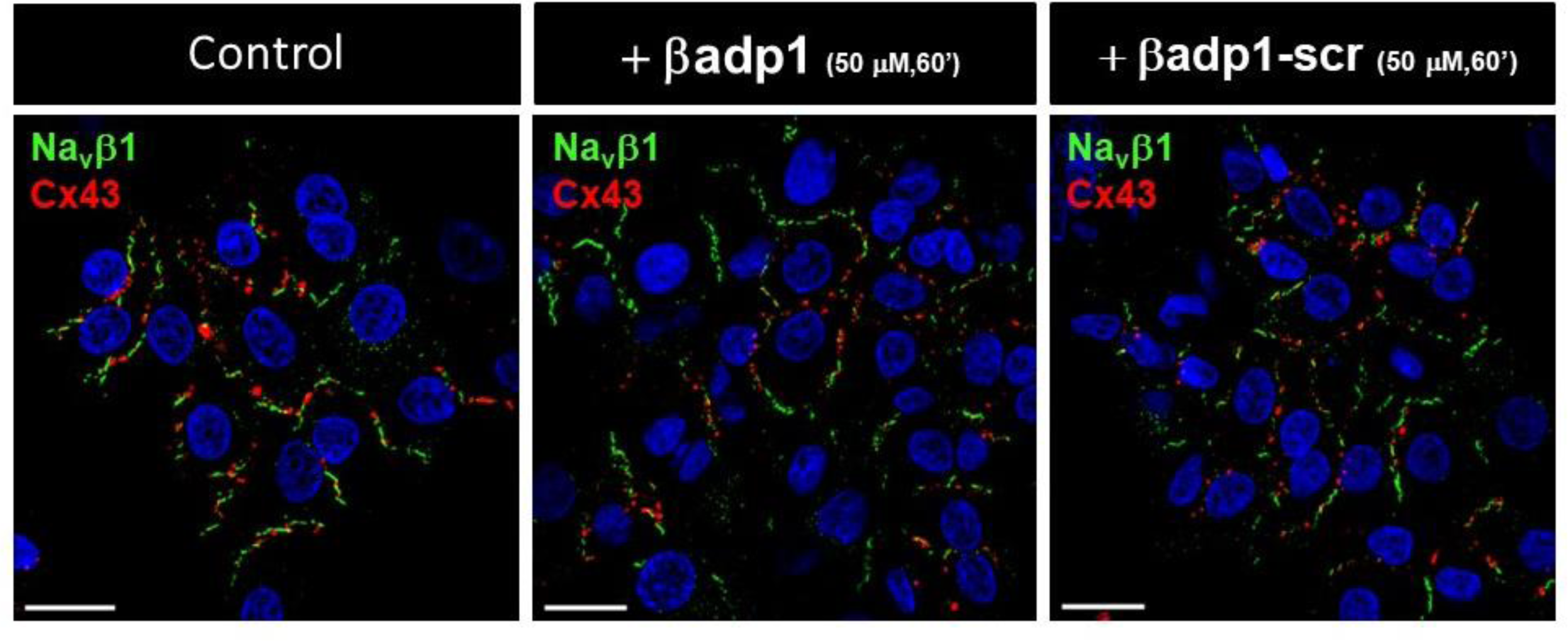
Confocal immunolabeling of NRVMs treated with scrambled control peptide for 60 minutes and immunolabeled for β1 Cx43: Representative confocal images of NRVMs immunolabeled for Cx43 (red) and β1 (green) of control NRVMs or after treatment with βadp1 or scrambled βadp1 control peptide at 50 μM for 60 minutes. Quantification of scrambled βadp1 control peptide is included in Fig 1. and shows no significant differences in junctional density of SCN1B (β1/β1B) compared to control NRVMs.

**Supplemental Figure 2.**
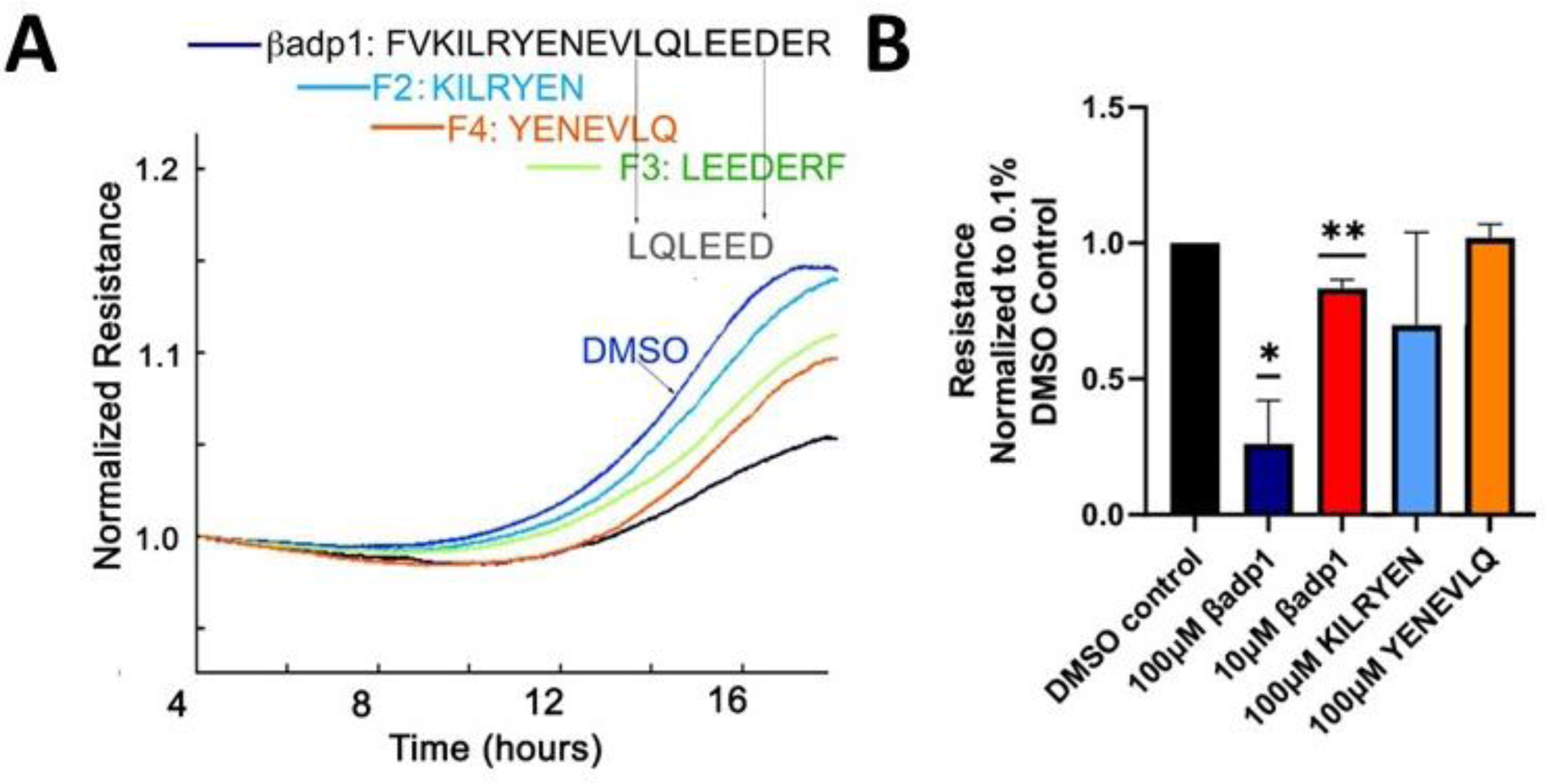
Effects of βadp1 and short βadp1-based monomeric sequences on relative resistance in 1610β1 cells βadp1: **A and B)** Multi-well ECIS demonstrating activity of βadp1 and short 6-8 amino acid N-terminal truncation variant peptides of βadp1. βadp1 (black) reduces normalized resistance in these cells relative to vehicle control (blue) over 20 hours. Peptides towards the carboxyl terminus of βadp1, especially those incorporating hydrophobic leucine residues, show the most adhesion reducing activity. Based on the ECIS time course data, and predictions from molecular modeling of likely binding in the hydrophobic pocket β1/SCN1B Ig domain binding surface, the sequence LQLEED was selected as the most effective short peptide candidate drug molecule. n=3 per peptide, *p<0.05, **p<0.01.

**Supplemental Figure 3.**
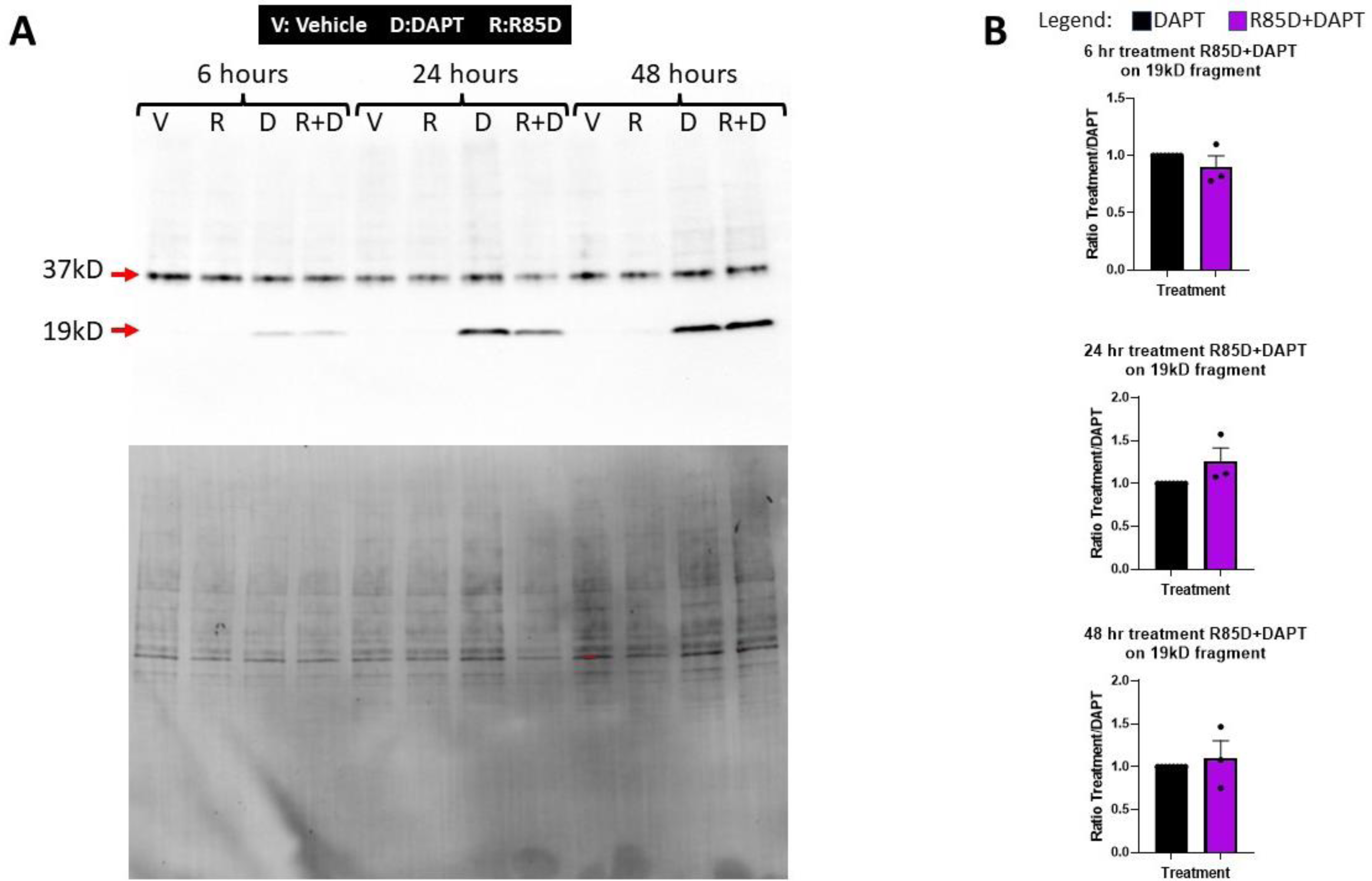
βadp1 control peptide R85D shows no effects on β1 RIP: **A and B)** Western blot analysis demonstrating R85D effects on the 19 kD β1 CTF generated by RIP. Blot shows effects following treatment with vehicle control (DMEM/F12 culture media with 0.1% DMSO), 1 μM DAPT, 50μM R85D, 50μM R85D +1μM DAPT at 6, 24 and 48 hours of treatment. Below the blot in (**A)** is the total protein control blot. **B)** Quantification of the β1 CTF at 6, 24, and 48 hours normalized to vehicle control, indicating no significant difference in the 19kD fragment when Scr1 treatment is present together with DAPT, relative to DAPT alone at the 6, 24 and 48 hours of a two-day treatment time course. n=3 experimental replicates for each treatment.,

